# Machine learning-based rescoring with MS²Rescore boosts peptide identification and taxonomic specificity in metaproteomics

**DOI:** 10.1101/2025.02.17.638783

**Authors:** Xuxa Malliet, Arthur Declercq, Ralf Gabriels, Tanja Holstein, Bart Mesuere, Thilo Muth, Pieter Verschaffelt, Lennart Martens, Tim Van Den Bossche

## Abstract

**Background:** Metaproteomics, the study of the collective proteome within microbial ecosystems, has gained increasing interest over the past decade. However, peptide identification rates in metaproteomics remain low compared to single-species proteomics. A key challenge is the identification sensitivity of current identification algorithms, which were primarily designed for single-species analyses. Addressing this, we evaluated the machine learning-driven MS²Rescore post-processing tool on multiple metaproteomics datasets from diverse microbial environments and benchmark studies.

**Results:** We demonstrate that machine learning-driven rescoring outperforms traditional metaproteomics identification workflows. It significantly increases peptide identification rates compared to Sage, which itself already implements basic rescoring. Moreover, it enables lowering the false discovery rate (FDR) to 0.1% with minimal to no sensitivity loss, a substantial improvement over the 1% or 5% FDR thresholds commonly used in metaproteomics, in turn leading to greater confidence in downstream taxonomic annotation.

**Conclusions:** Our findings show that MS²Rescore substantially improves peptide identification sensitivity as well as specificity in metaproteomics, and delivers improved confidence in taxonomic annotation. This advancement results in a more reliable downstream taxonomic analysis, reinforcing the potential of machine learning-based rescoring in metaproteomics research.

## Background

Metaproteomics, the study of the collective proteome within microbial ecosystems, complements metagenomics and metatranscriptomics by providing direct functional insights into complex microbial communities. It has been applied across diverse environments, including the human gut (1), biogas plants (2), and soil (3). However, the field continues to suffer from lower identification rates compared to single-species proteomics. The primary challenge lies in the limited sensitivity and statistical validation of current identification algorithms, which were originally designed for single-species datasets (4,5). In metaproteomics, analyses rely on large and highly diverse protein sequence databases. When applying the target-decoy approach for false discovery rate (FDR) control, the expanded search space increases the frequency of high-scoring decoy matches. Therefore, to maintain a desired estimated FDR (e.g. 1%), a high PSM score threshold is needed, which results in a loss of confidently accepted true identifications, as illustrated in **Figure 1**. Additionally, as datasets increase in complexity, taxonomic annotation becomes more difficult due to sequence homology issues (6). This necessitates a more refined peptide-level analysis to improve sensitivity and specificity.

**Figure 1.**
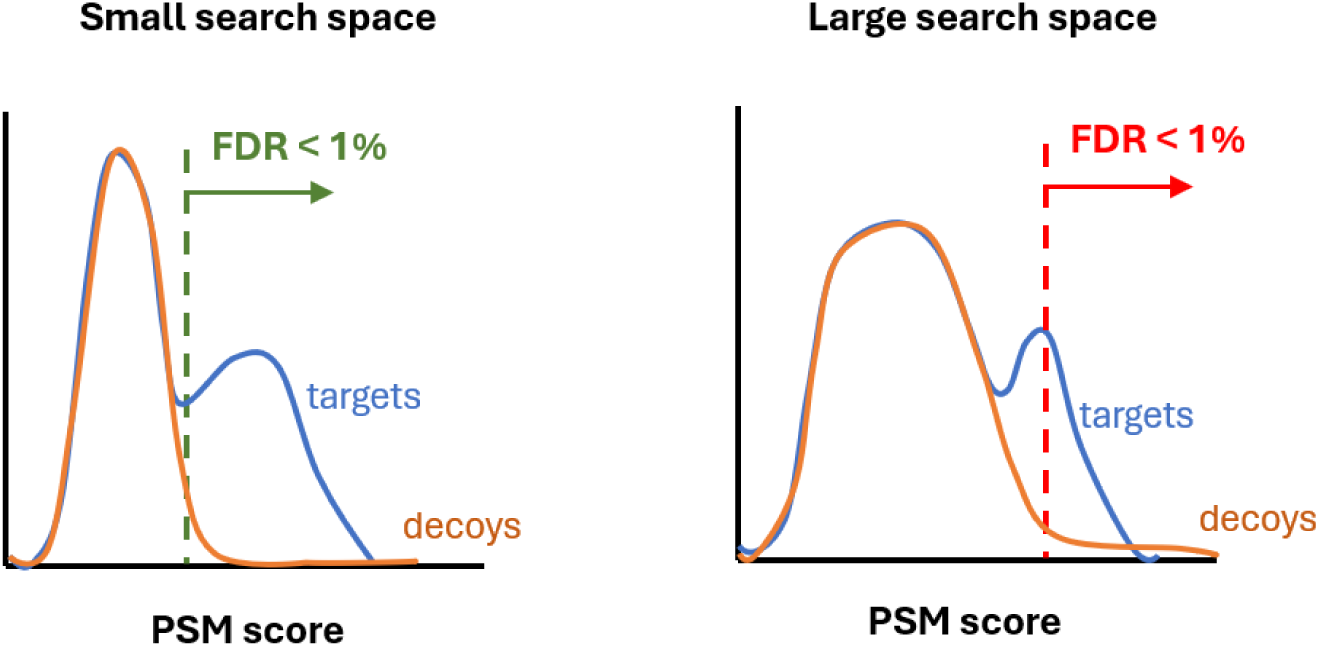
Impact of search space size on PSM score distributions. In smaller databases, decoy scores (orange) are well-separated from true target matches (second peak of blue distribution), while false target matches (first peak of blue distribution) overlap closely with the decoy distribution, enabling effective separation of true from false matches (green dashed line). In large search spaces, the decoy score distribution shows greater overlap with true target matches, resulting in a higher required threshold to maintain the estimated FDR of 1% (red dashed line), and a subsequent loss of true identifications.

To address these challenges, we applied the in-house developed machine learning-based MS²Rescore algorithm (7) to multiple multi-species metaproteomics datasets. Rescoring is a well-established post-processing strategy to improve identification rates, by training a machine learning model to better discriminate between target and decoy matches. A widely used example is Percolator (8), which allows users to define the feature set, typically derived from features produced by the search engine. MS²Rescore enhances standard rescoring by combining search engine-derived features with machine learning-derived features such as MS2 peak intensity predictions from MS²PIP (9) and retention time predictions from DeepLC (10). This approach not only increases peptide-spectrum match (PSM) identification rates, as previously demonstrated (11,12), but also enables a more stringent FDR threshold of 0.1%, improving upon the commonly used 1% or 5% thresholds in metaproteomics.

Besides improving peptide identification rates, we also evaluated the impact of MS²Rescore on taxonomic annotation. MS²Rescore substantially increased both PSM counts and the number of unique peptides for true-positive species. This high sensitivity of MS²Rescore exposes the known limitations of the lowest common ancestor (LCA) approach for taxonomic annotation, and further encourages the use of more sophisticated statistical methods such as Peptonizer2000 (13). Doing this, MS²Rescore identifications yield higher confidence in taxon identification.

## Methods & Implementation

### Datasets

To evaluate MS²Rescore for metaproteomics, we conducted three experiments using benchmark, controlled-mixture, and large-scale real-world datasets.

First, we reanalyzed mass spectrometry files from the CAMPI study (PXD023217) (6), the first multi-laboratory benchmark study in metaproteomics. We compared all data acquired with Thermo Fisher Scientific instruments, hence excluded datasets F03, F08, F09, S13, and S14, which were acquired using Bruker instruments. However, MS²Rescore does support rescoring data from these instruments as shown in Declercq et al. (14). As in the CAMPI study, all samples were searched against both the reference database and the multi-omic database. This experiment assessed how MS²Rescore performs in terms of peptide identification rates compared to the Sage search engine without additional rescoring.

Second, we analyzed four raw files from the iPRG 2020 (Phase I) metaproteomics study (PXD034795) (15) to determine the taxonomic specificity of MS²Rescore. The four raw files resulted from four mixtures with identical composition (3 mL of *Bacillus subtilis*, 30 mL of *Salmonella enterica*, and 30 mL of T4-infected *Escherichia coli*). However, all searches were conducted under the assumption of a completely unknown composition.

Third, we reanalyzed publicly available datasets from PRIDE (16): an inflammatory bowel disease (IBD) dataset consisting of 77 raw files (3.3 million spectra) (PXD010371) (1), a biogas plant (BGP) dataset with 441 raw files (14.3 million spectra) (PXD009349) (2), and a soil dataset containing 29 raw files (1.4 million spectra) (PXD005447) (3). These datasets were selected to assess MS²Rescore’s performance across three commonly studied metaproteomics environments (human gut, biogas plants, and soil) and across different fragmentation methods, with trap-type collision induced dissociation (CID) for the IBD and BGP datasets, and beam-type CID, also known as higher-energy collisional dissociation (HCD), for the soil dataset.

### Data processing with Sage

All raw files were converted to the mzML format using ThermoRawFileParser v1.4.5 (17), and peptide identification searches were conducted with Sage (18). Sage is a modern, open-source proteomics search engine that applies an internal rescoring strategy conceptually similar to Percolator (8). In brief, Sage uses linear discriminant analysis to distinguish target from decoy matches based on its own search engine-derived features. These features also include retention-time prediction errors derived from a per-search linear regression model. Given the fact that Sage already includes basic rescoring, we did not compare with baseline rescoring without analyte prediction features.

Notably, Sage is exceptionally fast, while still matching the performance of other state-of-the-art search engines. Similarly to the MSFragger search engine (19), Sage achieves this speed partly through fragment indexing. Here, observed MS2 ions are mapped to all peptides in the database that could have produced this fragment. Keeping the full fragment ion index in memory, this approach enables very efficient searching but also implies that memory usage scales with database size. To address this limitation for very large metaproteomics databases, a beta release of Sage (v0.15.0-beta.1) featuring internal database splitting was used for all experiments. Exceptions were the SIHUMIx samples and the fecal samples searched against the multi-omic database (CAMPI study, first experiment), which were searched with Sage v0.14.7, as these did not require database splitting. While using internal database splitting does reduce the memory required for large database searches, it considerably reduces Sage’s normally high processing speed compared to when the full database is loaded directly, such as in v0.14.7.

### Search parameters

For the first and third experiments, peptide identification was performed against a concatenated target/decoy database including all sequences from the original studies (1–3,6). Since the composition of the iPRG 2020 samples was unknown (second experiment), we searched against UniRef50 (20) (39,232,797 sequences). The database had to include sequences from all domains (Archaea, Bacteria, and Eukarya), as well as from viruses and prions. The UniRef50 database provides clustered sets of sequences from the UniProt Knowledgebase (including isoforms) and selected UniParc records in order to obtain as complete as possible a coverage of the sequence space at several resolutions, while hiding redundant sequences. To ensure all contaminant proteins were included, the UniRef50 database was concatenated with the common repository of adventitious proteins (cRAP) database (116 sequences, https://www.thegpm.org/crap/).

For all databases, decoy sequences were generated by reversing target sequences. Identification settings were identical to those described in the original manuscripts: cleavage with trypsin (C-terminal of lysine or arginine, except when followed by proline) with a maximum of two missed cleavages (all datasets); 10.0 ppm as MS1 tolerance (all datasets); 0.02 Da (soil, CAMPI), 0.5 Da (IBD, BGP), or 0.6 Da (iPRG) as MS2 tolerance; Carbamidomethylation of C as fixed modification (all datasets); Oxidation of M as variable modification (all datasets). Sage has the option to exclude certain fragments from the fragment index, which helps to reduce memory usage. Here, we excluded b1/b2/y1/y2 ions by setting min_ion_index to 2, as they are frequently shared across peptides, and needlessly increase the fragment ion index while slowing down searching. The output TSV files from Sage were used for post-processing with MS²Rescore.

### MS²Rescore post-processing

Here, we used MS²Rescore v3.2.0.post1 for rescoring the identifications produced by Sage. MS²Rescore allows the user to decide whether Percolator or Mokapot is used as an internal rescoring engine. Here, Mokapot was selected because it integrates more efficiently with MS²Rescore.

MS²Rescore is available as a Python package (https://github.com/compomics/MS2Rescore) and includes a Graphical User Interface (GUI). The tool is open-source under the Apache 2.0 license.

### Data analysis

For the first experiment, unique peptides identified by MS²Rescore and the search engines used in the CAMPI study were visualized using UpSet plots. Peptides were compared without modifications, with leucine (L) and isoleucine (I) considered equivalent (converted to J). Peptide identifications from the CAMPI study were retrieved from Supplementary Data 15 and 16 (6).

For the second experiment, taxonomic analysis was performed using the Unipept CLI v4.0.1 (https://unipept.ugent.be) (21–23), with the *pept2lca* function, and Peptonizer2000, used here as a standalone tool (available at https://github.com/compomics/Peptonizer2000), though it is also available through Unipept (13).

For the third experiment, we compared the identification rates at different FDR thresholds for all three analyzed public datasets.

### Data availability

All .mzML files, configuration files and results files from Sage and MS2Rescore are available via Zenodo (https://doi.org/10.5281/zenodo.18757265), as well as all Jupyter Notebooks used to generate the figures presented in this manuscript. Source code for MS²Rescore can be consulted via GitHub (https://github.com/compomics/MS2Rescore).

## Results and Discussion

### MS²Rescore improves peptide identification in metaproteomics

In this first experiment, we analyzed all samples using Sage and Sage + MS²Rescore, then calculated the improvement in identification rates after rescoring at 0.1% and 1% FDR thresholds. Identification rate is defined as the proportion of acquired spectra that were assigned a peptide identification, and **Figure 2** shows the relative percentage increase upon rescoring. Across all samples, MS²Rescore increased the identification rate, with all improvements being higher than zero. The largest gains were observed in the more complex conditions: searches against the REF database yielded higher improvements than MO, and fecal samples showed greater increases than SIHUMIx samples. Additionally, improvements were more pronounced at the 0.1% FDR threshold compared to the 1% FDR threshold.

**Figure 2.**
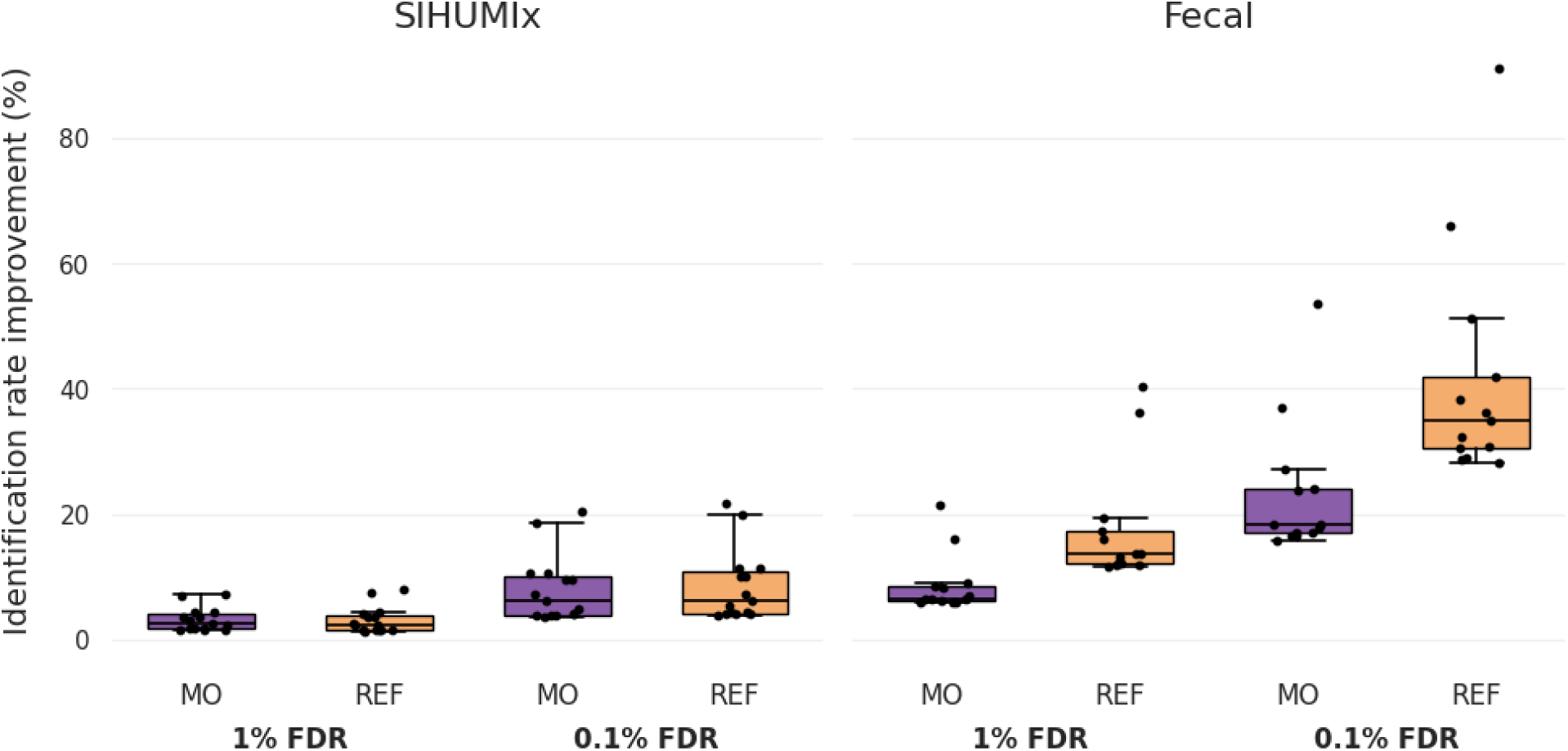
Boxplots showing the improvement in identification rates (relative percentage increase) after rescoring Sage results with MS²Rescore for all CAMPI study samples. Results are grouped by sample type (SIHUMIx, left; fecal, right), search database (MO in purple, REF in orange), and FDR threshold (1% and 0.1%).

The primary objective of this experiment was to benchmark MS²Rescore against widely used search engines in metaproteomics, as described in the CAMPI study (6). These pipelines include MetaProteomeAnalyzer (X!Tandem and OMSSA), Proteome Discoverer (SequestHT), MaxQuant (Andromeda), and SearchGUI/PeptideShaker (X!Tandem, OMSSA, MS-GF+, and Comet). **Figure 3** shows an upset plot comparing all these pipelines to the Sage + MS²Rescore workflow for the F07 sample searched against the MO database. An equivalent comparison for the S11 sample searched against the REF database is shown in **Supplementary Figure 1**.

**Figure 3.**
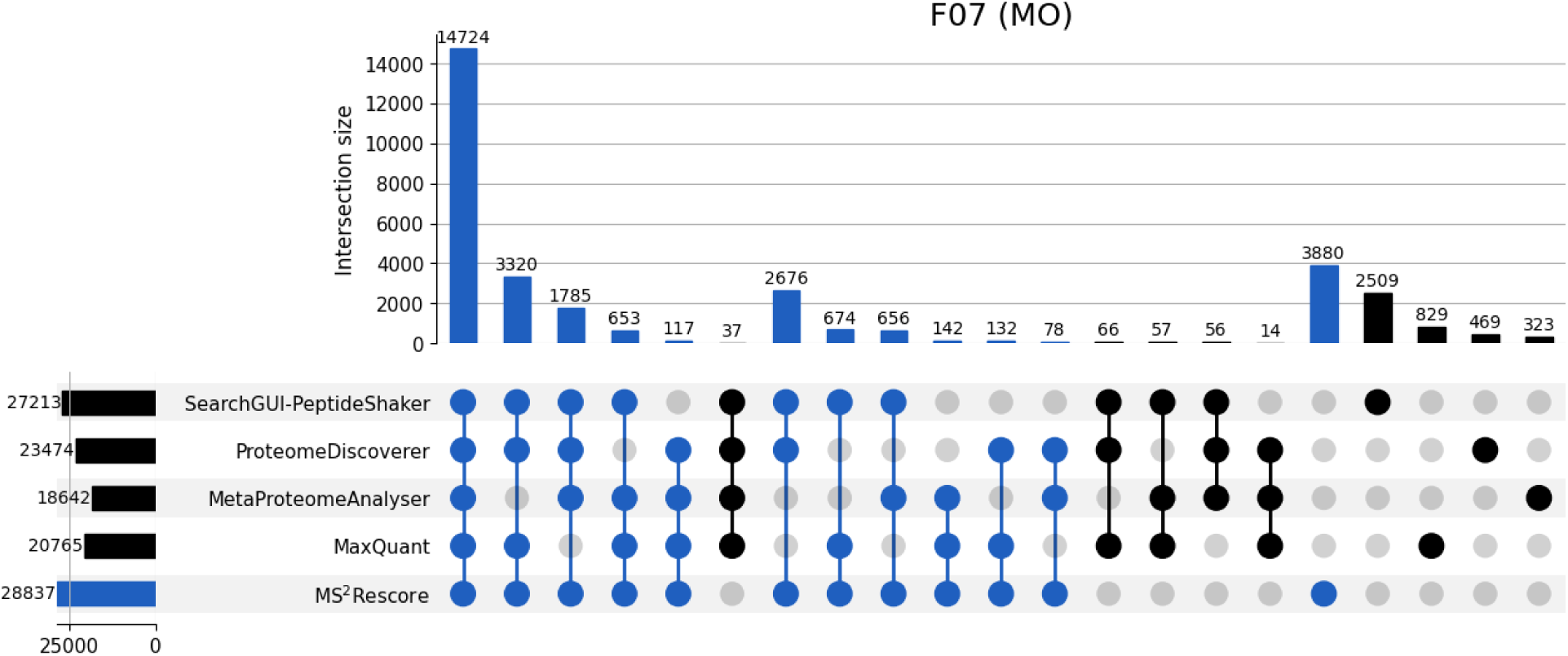
Comparison of Sage + MS²Rescore with the four original identification pipelines used in the CAMPI study for the fecal sample F07. The x-axis lists different identification pipelines: MetaProteomeAnalyzer (MPA, using X!Tandem and OMSSA), Proteome Discoverer (PD, using SequestHT), MaxQuant (MQ, using Andromeda), and SearchGUI/PeptideShaker (PS, using X!Tandem, OMSSA, MS-GF+, and Comet). Blue bars show overlapping peptides found by MS²Rescore, whereas black bars indicate peptides uniquely identified by other pipelines. Sets with just two pipelines were excluded for visualization purposes.

In total, 33,197 peptides were identified for the fecal F07 sample, of which 44.4% were detected by all identification pipelines. Interestingly, MS²Rescore identified 86.9% of all identified peptides.

It is also noteworthy that MS²Rescore outperformed the best-performing pipeline from the CAMPI study. In the original study, SearchGUI/PeptideShaker yielded the highest number of unique peptide identifications by integrating results from four search engines (X!Tandem, OMSSA, MS-GF+, and Comet). However, our results demonstrate that MS²Rescore further increases peptide identifications, reinforcing prior suggestions from the CAMPI study that machine learning-based search engines could significantly enhance metaproteomics identification rates. Moreover, this means that comparable or better performance can be obtained without running and harmonizing four separate search engines. Importantly, MS²Rescore still recovers the majority of peptides that were consistently identified across the CAMPI search workflows, indicating little to no loss of putatively correct identifications. Furthermore, among the unique identifications not retained by MS²Rescore, 80% were supported by only a single search engine in the original analysis, suggesting that these are likely spurious.

### Higher confidence in identifications improves taxonomic annotation confidence

In this second experiment, we evaluated the impact of MS²Rescore on taxonomic annotation using data from the iPRG 2020 study. First, we compared the number of confidently identified spectra across different FDR thresholds for Sage and Sage + MS²Rescore. **Figure 4** shows highly similar q-value plots for all four lab-generated mixes, indicating highly reproducible identification sensitivity and specificity.

**Figure 4.**
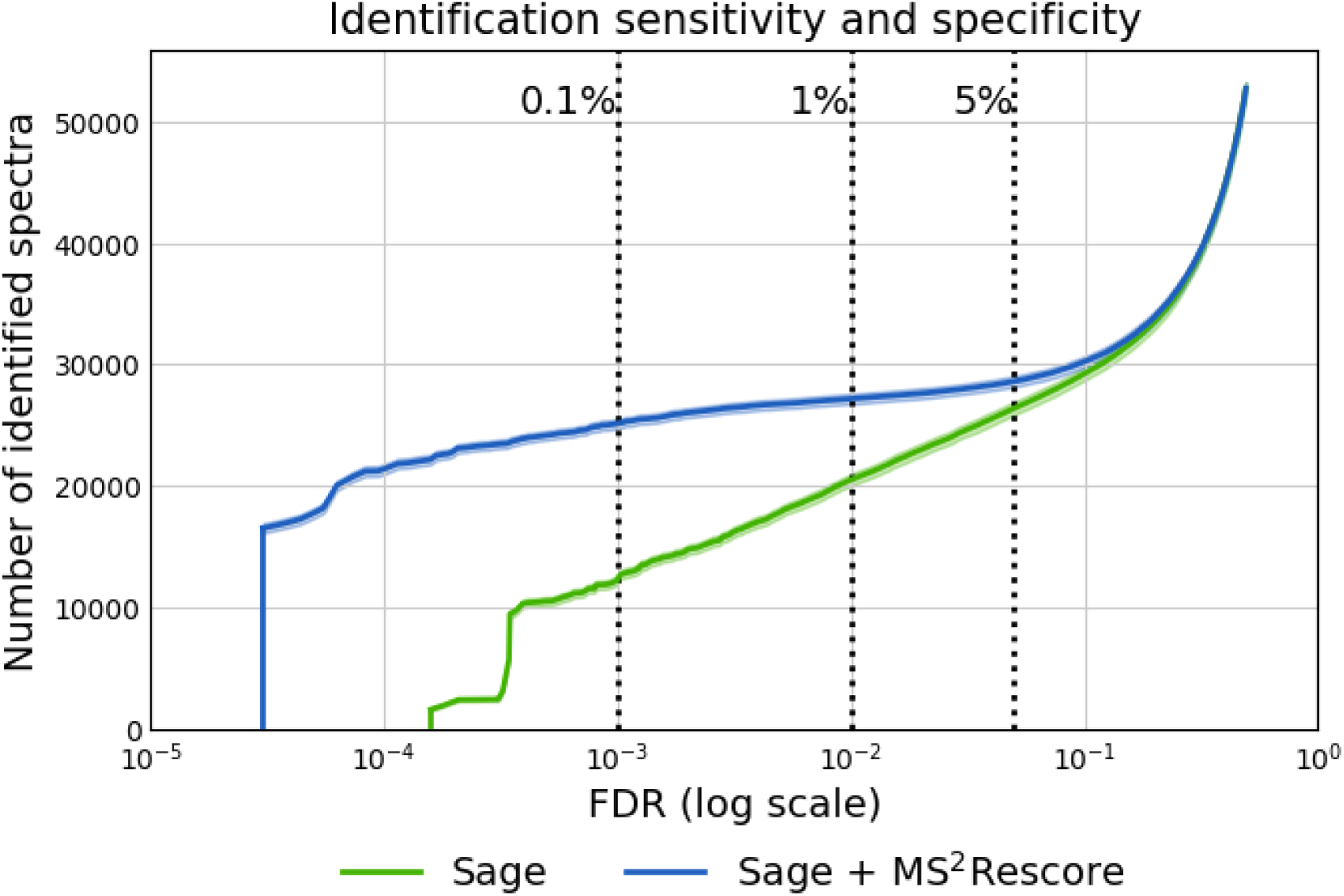
Number of identified spectra across varying false discovery rate (FDR) thresholds for Sage ( green) and Sage plus MS²Rescore (blue). Averages across the four mixes are plotted, with variation between mixes indicated by the shaded areas. Vertical dotted lines indicate FDR thresholds of 0.1%, 1%, and 5%.

Sage + MS²Rescore consistently outperformed standalone Sage in terms of peptide identification rates, yielding about twice as many identifications at an FDR of 0.1%. Importantly, rescoring with MS²Rescore therefore allows the estimated FDR to be reduced from 1% to 0.1% with minimal loss of identifications.

To illustrate the effect of improved peptide identification on downstream taxonomic annotation, we analyzed the PSMs identified by MS²Rescore in Mix 1 at both 1% and 0.1% FDR thresholds using the online Unipept server to obtain a treeview for both. At 1% FDR, a high number of unexpected and biologically implausible taxa were detected (**Figure 5**), suggesting that many assignments were false positives. Conversely, at a 10-fold higher specificity of 0.1% FDR, a much more distinct set of taxa were detected. This shows the benefit of lowering the FDR threshold from 1% to 0.1% for taxonomic annotation.

**Figure 5.**
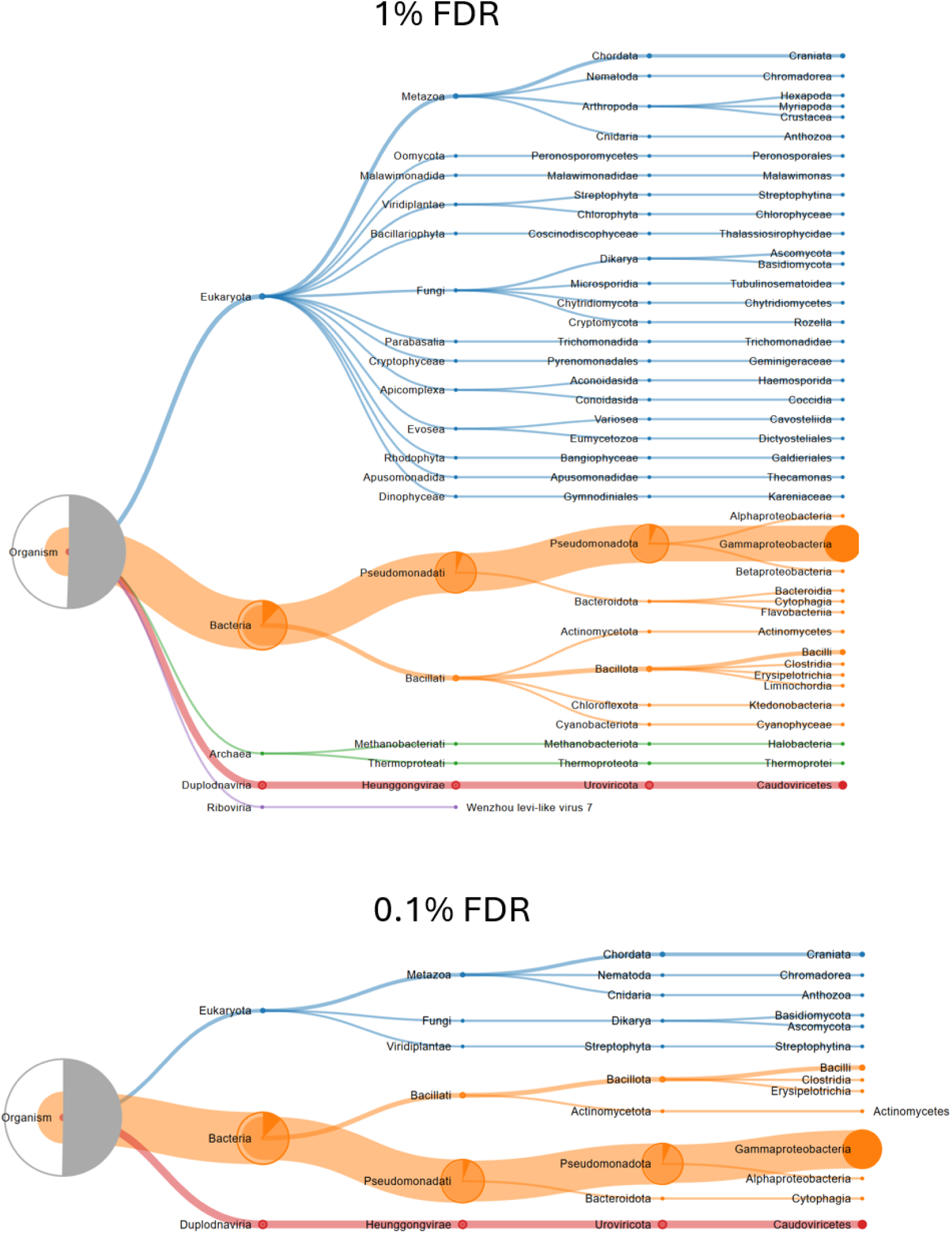
Unipept taxonomic annotations after MS²Rescore identification at different confidence levels. The top panel shows the Unipept treeview for Mix 1 at 1% false discovery rate (FDR), while the bottom panel represents Mix 1 at 0.1% FDR. These treeviews are presented for illustrative purposes, demonstrating how a stricter FDR threshold reduces the number of identified taxa, improving specificity while maintaining biologically relevant taxa.

For the actual taxonomic analysis, we first applied a straightforward approach using Unipept’s LCA method. PSMs from the four mixtures were filtered at a stringent 0.1% FDR, and peptides matching any proteins derived from the cRAP database were removed. The *pept2lca* function was used to retrieve the LCA for all peptides in each mixture individually. Species-level LCAs detected in at least three of the four mixtures were retained. For these, the total number of PSMs corresponding to their peptides was computed. In **Figure 6**, bar heights represent the average spectrum count across the four mixtures, and numbers on top of the bars indicate the number of distinct peptides contributing to this count. The same analysis was performed for both Sage and Sage + MS²Rescore results.

**Figure 6.**
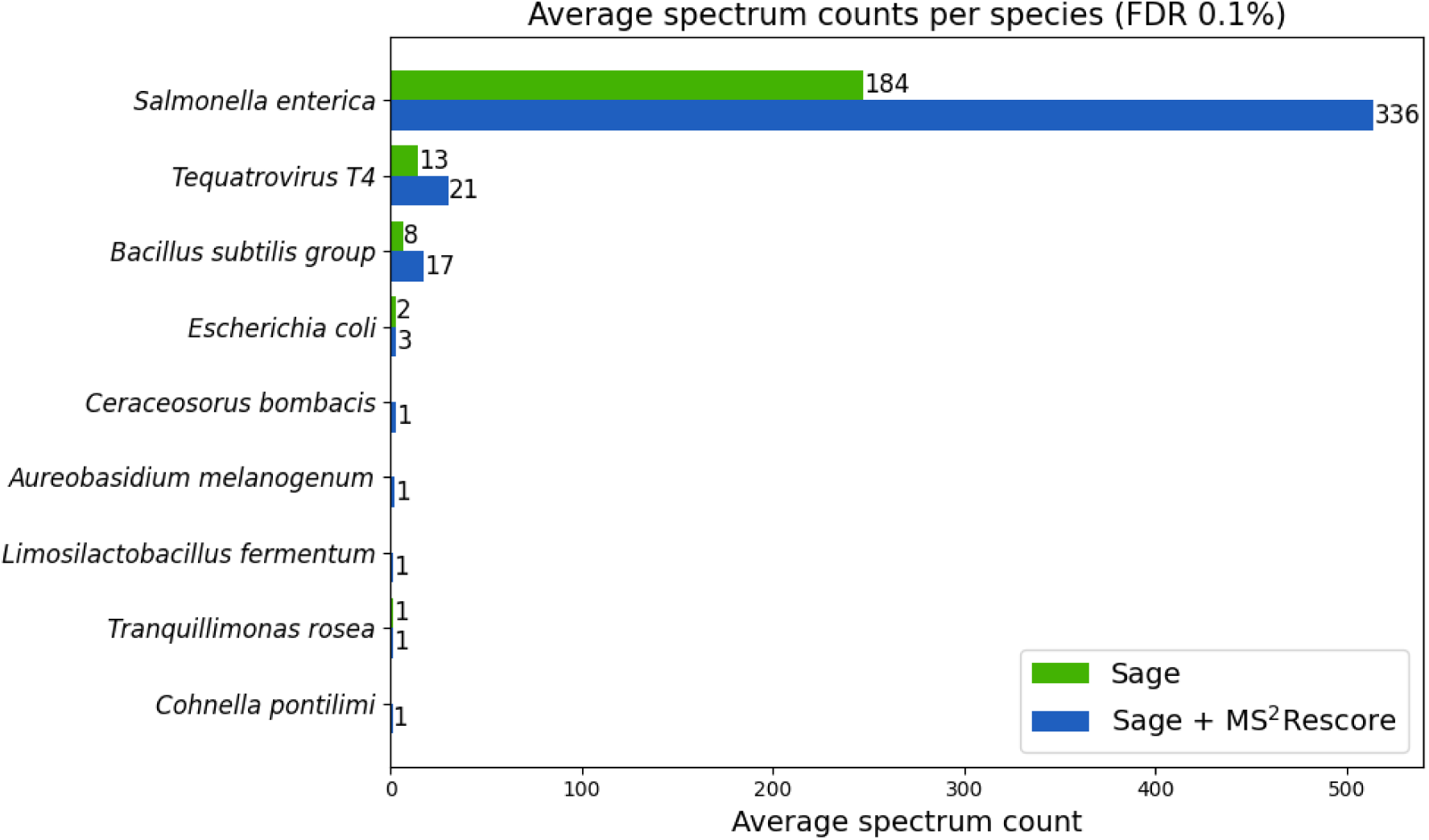
Bar plot of average spectrum counts across the four mixes for all species that occurred as a lowest common ancestor in at least 3 mixes, at a false discovery rate threshold of 0.1%. Bar height shows average spectrum count, while numbers on top of the bars indicate the number of unique peptides responsible for these spectrum counts. Sage is shown in green, Sage + MS²Rescore is shown in blue.

Using this simple LCA-based approach, both Sage and Sage + MS²Rescore successfully identified all four species present in the mixtures. Leveraging MS²Rescore yielded more unique peptides per species and higher overall spectrum counts. Notably, *Escherichia coli* showed substantially lower species-level spectrum counts compared to *Salmonella enterica*, despite equal sample abundance. This discrepancy arises because most *E. coli* peptides are shared with other species, resulting in very few peptides with *E. coli* as the LCA and, consequently, few PSMs assigned at the species level. Applying Unipept’s *pept2taxa* function confirmed that *E. coli* had the highest number of matching peptides, but nearly all had LCAs above the species level, indicating they are shared with other species (**Supplementary Figure 2**). Van Den Bossche et al. (24) reported that this issue can be largely avoided using a custom Unipept database; however, this was not possible in the present study due to the assumption of unknown sample composition.

While LCA-based analysis identified the target species, it also revealed a small number of apparent false positives. Sage + MS²Rescore identified five additional false-positive species, whereas Sage identified only one. At first sight, this suggests that although MS²Rescore yields a substantial increase in true-positive peptide identifications, it is accompanied by a small increase in false positives at the taxonomic level. However, a closer examination showed that many of these species-level false positives are likely artifacts of preliminary or non-curated database entries rather than true misidentifications. For example, peptides mapping to *Ceraceosorus bombacis*, *Aureobasidium melanogenum*, *Limosilactobacillus fermentum*, *Tranquillimonas rosea*, and *Cohnella pontilimi* are either derived from WGS datasets, homology-based predictions, or non-reference proteomes. BLAST searches confirmed that several of these peptides correspond to hypothetical proteins. Further details are provided in the **Supplementary Information**.

Beyond database issues, the fundamental limitation arises from the LCA approach itself. LCA discards all peptides shared between taxa, even when they provide strong cumulative evidence for species presence. As a result, genuine peptide evidence can be lost, as observed for *E. coli*, and rare falsely identified peptides (which are still expected at a 0.1% FDR threshold) can have a disproportionate effect on the taxonomic results. Indeed, a single falsely identified peptide can lead to an entire falsely identified species. The latter effect becomes more visible when using MS²Rescore simply because the higher sensitivity results in substantially more identified peptides, and therefore also a higher absolute number of false identifications.

To overcome these issues, we applied Peptonizer2000, which integrates all peptides through a statistical framework that accounts for sharing and identification confidence. This probabilistic approach limits the impact of single false-positive peptides on species-level inference, producing scores that reflect confidence in species presence. Applying Peptonizer2000 to the MS²Rescore results, all top-scoring species correspond to actual sample composition, including *E. coli*, while falsely identified species receive substantially lower confidence (**Figure 7**).

**Figure 7.**
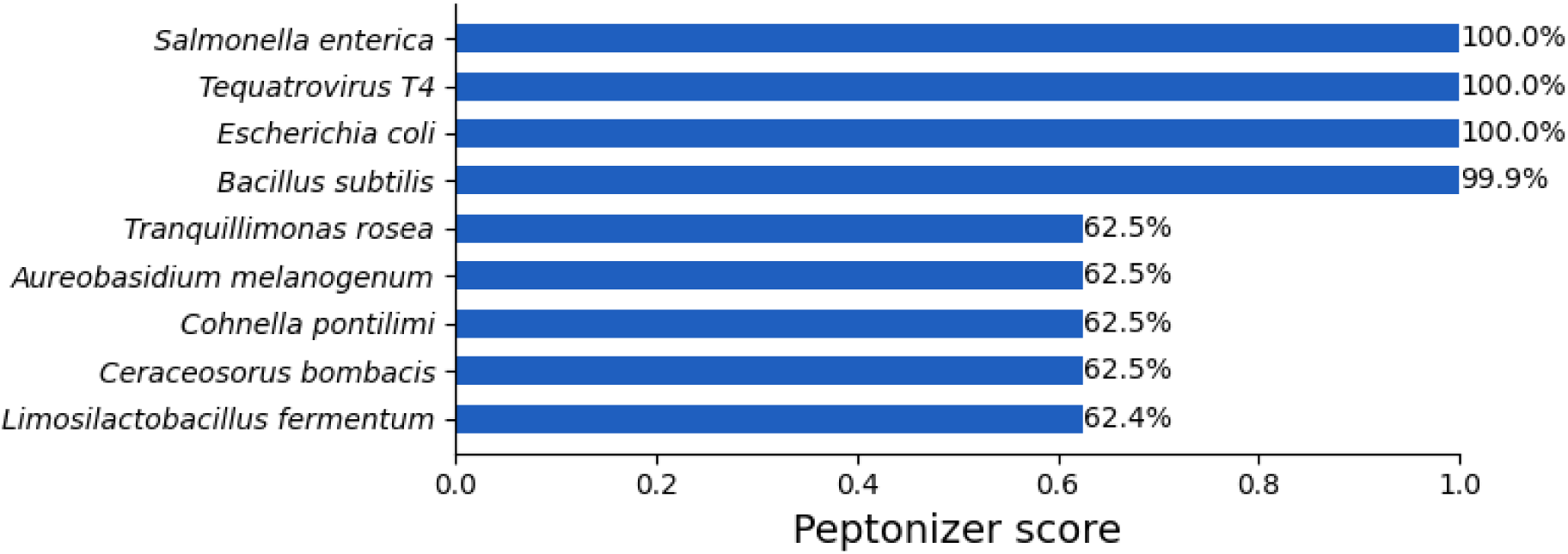
Peptonizer2000 confidence scores based on MS²Rescore peptide identifications for species present in the sample (top four) and species that were falsely identified using the LCA-based approach (bottom five).

Furthermore, applying Peptonizer2000 to the Sage results alongside the MS²Rescore results demonstrates that non-present species receive lower confidence scores with MS²Rescore, resulting in a shorter, more accurate species list (**Supplementary Figure 3**). This demonstrates that Peptonizer benefits from the higher sensitivity of MS²Rescore, which improves both confidence and specificity. To translate these performance gains into a usable resource, Peptonizer2000 was integrated into the Unipept web application in early 2025 (v6.1.0), offering a user-friendly and streamlined interface for high-confidence metaproteomics analysis.

In summary, while LCA provides a basic species-level overview, it is highly sensitive to peptide sharing and rare false positives. Leveraging MS²Rescore increases peptide identifications, and combining it with statistical frameworks like Peptonizer2000 produces robust species-level assignments, accurately reflecting sample composition and mitigating the limitations of traditional LCA-based analysis.

### MS²Rescore increases identification rates in large, publicly available metaproteomics datasets from diverse environments

In this third experiment, we further evaluated MS²Rescore’s performance in three large-scale studies, each representing one of three widely studied metaproteomics samples: human gut (IBD), biogas plant (BGP), and soil. **Figure 8** shows the identification rates at different FDR levels for each dataset.

**Figure 8.**
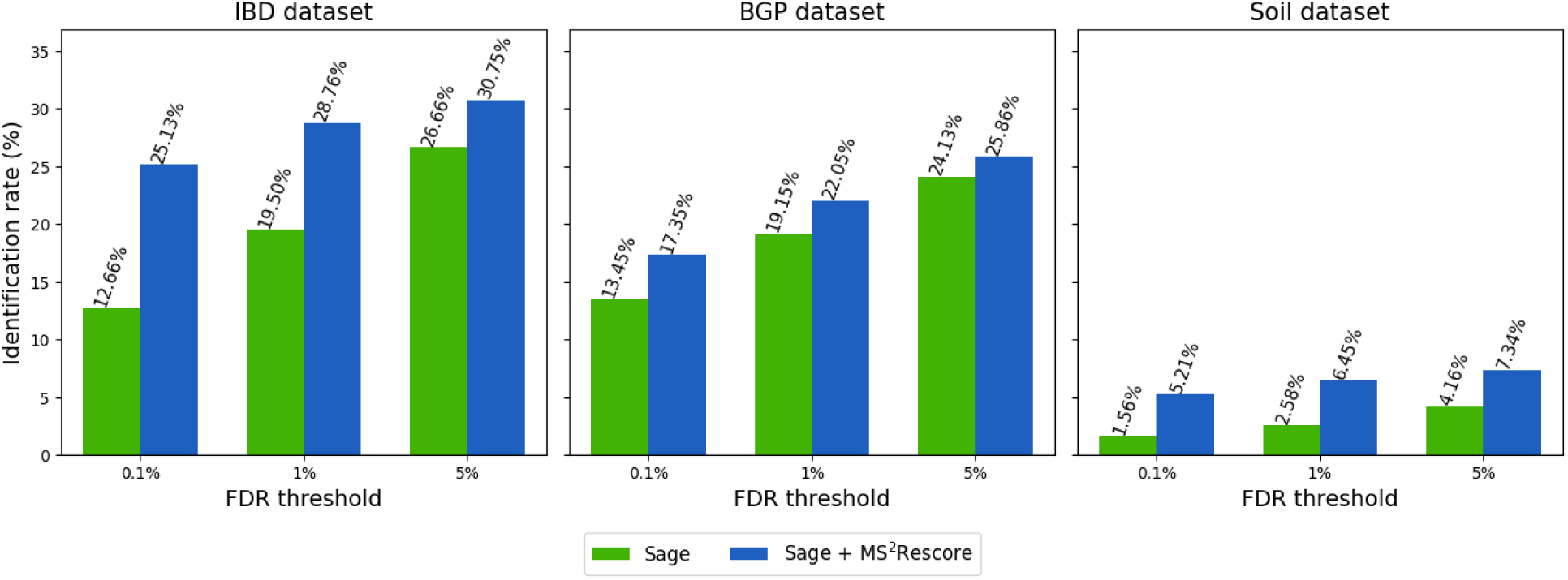
Identification rates for the IBD, BGP, and soil metaproteomics datasets at varying FDR thresholds (0.1%, 1%, and 5%) for Sage (green) and Sage + MS²Rescore (blue).

Across all datasets and thresholds, Sage + MS²Rescore outperforms standalone Sage, with the largest increases at the 0.1% level. Especially noteworthy is MS²Rescore’s performance in the soil dataset, which is a notoriously complex environment for metaproteomics analysis. Here, the identification rate at 0.1% FDR for Sage + MS²Rescore is higher than the identification rate of standalone Sage at 5% FDR. These results show that MS²Rescore can be extremely helpful in complex metaproteomics challenges, even compared to a search workflow that already includes basic rescoring.

## Conclusion

In conclusion, MS²Rescore substantially improves peptide identification rates in metaproteomics, directly addressing a major limitation of the field. Across all datasets, this gain enabled the use of a more stringent 0.1% FDR threshold without loss of sensitivity. Its strong performance in metaproteomics stems from its ability to counteract the convergence of target and decoy score distributions that occurs by chance in large, inflated search spaces. By combining conventional search engine-derived features with additional predicted features, it enhances the separation between true and false PSMs and mitigates the adverse effects of database inflation. The resulting increase in both the number and confidence of peptide identifications markedly improves the reliability of downstream taxonomic analyses, often yielding approximately twice as many unique peptides per present species. Moreover, when combined with advanced statistical frameworks such as Peptonizer2000, these high-confidence peptide identifications can be integrated to produce robust species-level assignments, effectively mitigating the impact of shared or low-confidence peptides and reducing false-positive taxonomic calls. Based on these results, we strongly recommend the combined use of data-driven rescoring, a stringent estimated FDR of 0.1%, and statistical inference approaches like Peptonizer2000 to achieve accurate and interpretable metaproteomics analyses.

## Acknowledgements

X.M., A.D., R.G., L.M., and T.V.D.B acknowledge funding from the Research Foundation Flanders (FWO) [1SA6I26N, 3S004321, 12AK526N, G010023N, 1286824N]. P.V. acknowledges funding by Ghent University [BOF/01P10623]. L.M. acknowledges funding by the Ghent University Concerted Research Action (BOF21/GOA/033), and by the European Union’s Horizon Europe Programme [101080544, 101103253, 101195186, and 10119173].

## Supplementary figures

**Supplementary Figure 1.**
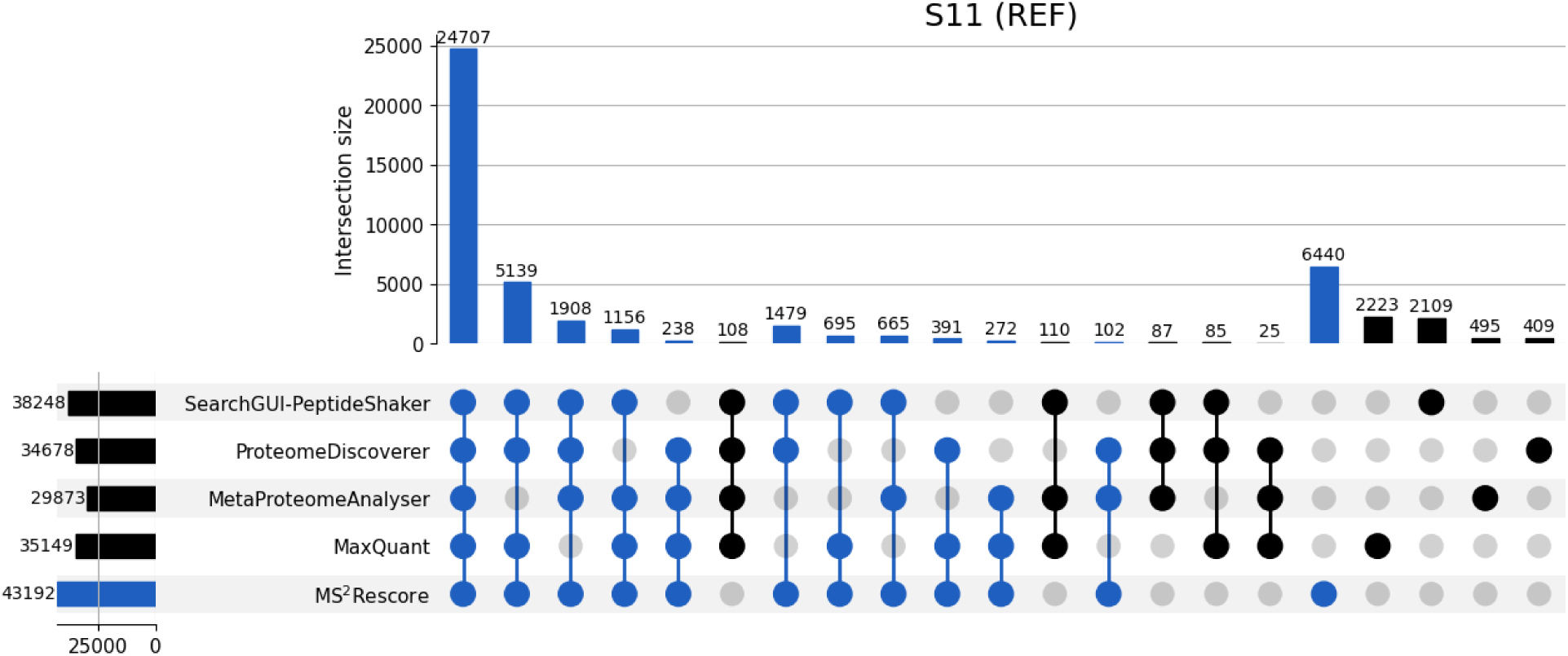
Comparison of Sage + MS²Rescore with the four original identification pipelines used in the CAMPI study for the SIHUMIx sample S11. The x-axis lists different identification pipelines: MetaProteomeAnalyzer (MPA, using X!Tandem and OMSSA), Proteome Discoverer (PD, using SequestHT), MaxQuant (MQ, using Andromeda), and SearchGUI/PeptideShaker (PS, using X!Tandem, OMSSA, MS-GF+, and Comet). Blue bars show overlapping peptides found by MS²Rescore, whereas black bars indicate peptides uniquely identified by other pipelines. Sets with just two pipelines were excluded for visualization purposes.

**Supplementary Figure 2.**
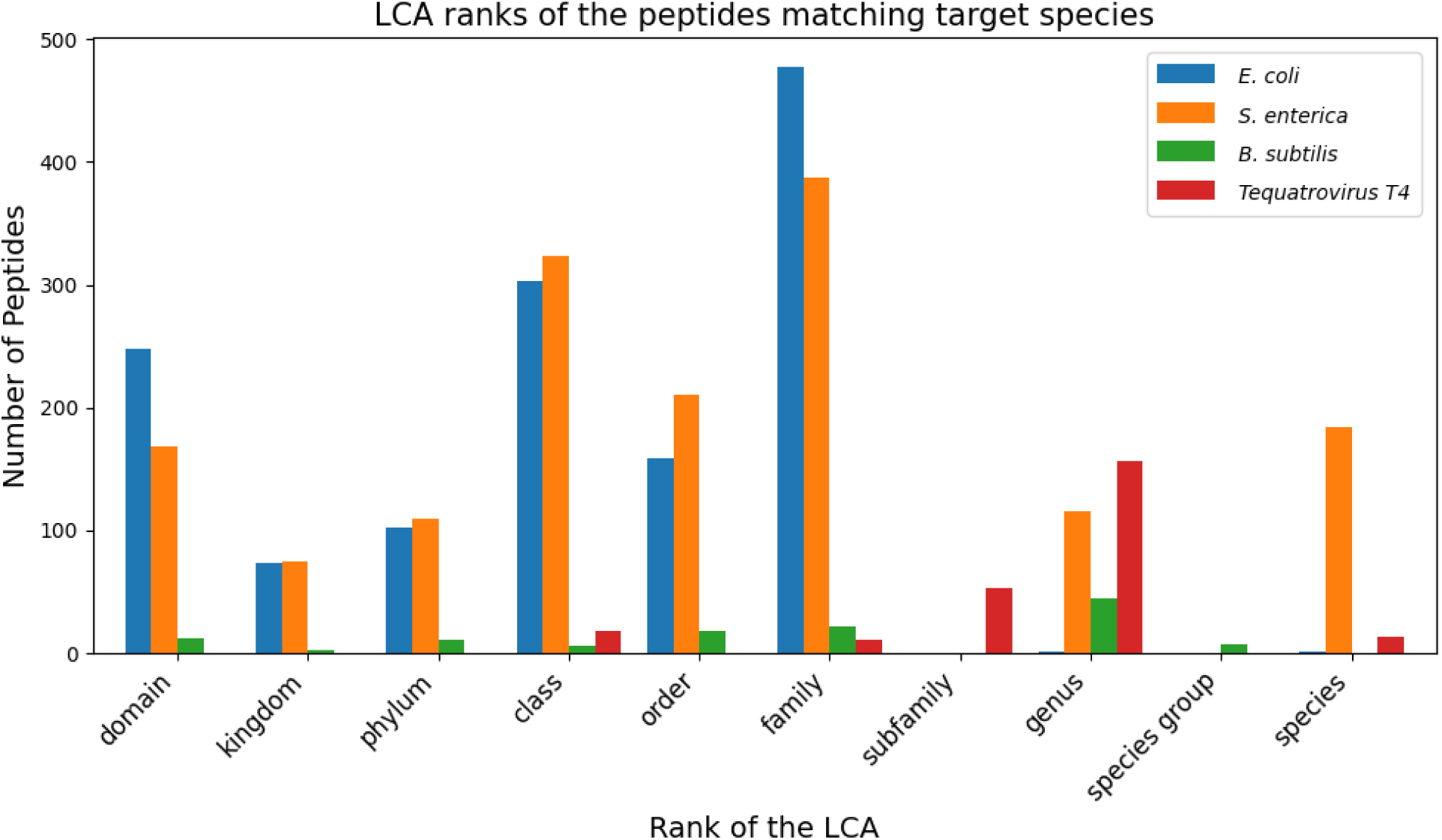
Ranks of the LCAs of all peptides matching one of the target species (*E. coli, S. enteria, B. subtilis, Tequatrovirus T4*) with Unipept *pept2taxa*. Many peptides matched *E. coli*, but almost all of them are shared, resulting in a higher-level LCA, and no identification of *E. coli* at species level.

**Supplementary Figure 3.**
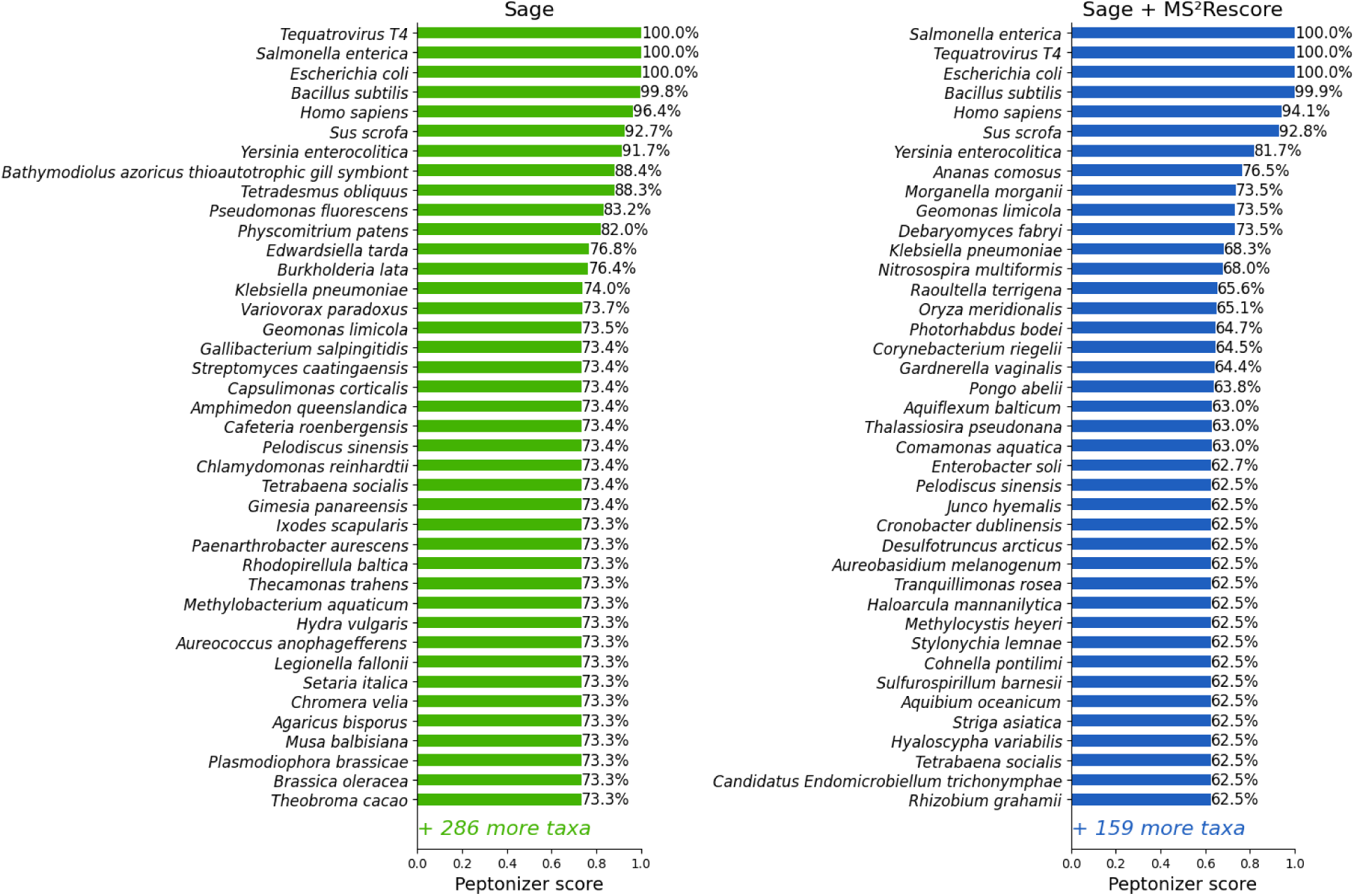
Peptonizer2000 confidence scores for all species after peptide identification with Sage (green) and Sage + MS²Rescore (blue). All top-scoring taxa correspond to species actually present in the sample. The first species not present (*Yersinia enterocolitica*) receives a confidence score of 91.7% with Sage, which decreases to 81.7% with Sage + MS²Rescore. This pattern continues for all lower-confidence species. Fewer species are reported in the MS²Rescore results, as Peptonizer only reports taxa with at least 50% confidence.

## Supplementary information

In the second experiment on taxonomic analysis, we investigated the five peptides that corresponded to the five false positively identified species in detail. Below is what we found for each, supporting our conclusion that these species-level false positives are likely artifacts of preliminary or non-curated database entries rather than true misidentifications.

The peptide IVSWYDNESGYSNR maps to the UniProtKB/TrEMBL entry IA0A0P1BR14 from Ceraceosorus bombacis. This protein, Glyceraldehyde-3-phosphate dehydrogenase, is inferred from homology and is widely present in all living organisms.

The peptide ERLELAEQHR maps to four UniProtKB/TrEMBL entries (A0A9P8J329, A0A074VNS6, A0A9P8FZR2, A0A9P8FTK7) from *Aureobasidium melanogenum*, all annotated as HTH APSES-type domain-containing proteins. Two of these entries are scheduled for removal from UniProtKB/TrEMBL because they are not part of a reference proteome, while the remaining two are predicted proteins derived from EMBL/GenBank/DDBJ whole genome shotgun (WGS) entries, which represent preliminary data. BLAST searches against NCBI nr further show that this peptide only maps to hypothetical proteins.

EAQQGLDQAK maps to five UniProtKB/TrEMBL entries (A0A829LVW3, A0A2K2TGJ6, D0DVV5, A0A843QX47, A0A1L7GSY7) from *Limosilactobacillus fermentum*, all of which are scheduled for removal from UniProtKB/TrEMBL because they are not part of any reference proteome. The peptide MSTIPVTLLTGFLGAGK, identified with both Sage and MS²Rescore, maps to a single UniProt entry (A0A1H9SW92), annotated as a GTPase (G3E family) from *Tranquillimonas rosea* inferred solely from homology.

EAAYLIGGLAALSR maps to A0A4U0F7R8, annotated as a glycosyltransferase RgtA/B/C/D-like domain-containing protein from *Cohnella pontilimi*, but this entry is also derived from a WGS dataset and represents preliminary annotation. These examples highlight that many apparent species-level false positives are likely the result of preliminary or non-curated database entries, rather than genuine misidentifications in the experimental data.

